# Lipid mediated inhibition of Niemann-Pick C1 protein is an evolutionary conserved feature of multiple *Mycobacterium* lineages and non-tubercular mycobacteria

**DOI:** 10.1101/2021.09.06.459187

**Authors:** Yuzhe Weng, Dawn Shepherd, Jiayun Zhu, Yi Liu, Nitya Krishnan, Brian D. Robertson, Nick Platt, Gerald Larrouy-Maumus, Frances M. Platt

**Affiliations:** Department of Pharmacology, University of Oxford, Mansfield Road, Oxford, OX1 3QT, UK; MRC Centre for Molecular Bacteriology and Infection, Department of Life Sciences, Faculty of Natural Sciences, Imperial College London, London, United Kingdom; MRC Centre for Molecular Bacteriology and Infection, Department of Infectious Disease, Flowers Building, Imperial College London, London, SW7 2AZ, UK

**Keywords:** *Mycobacterium tuberculosis*, Niemann-Pick Disease Type C, lysosome, lipid

## Abstract

*Mycobacterium tuberculosis* (*Mtb)* is the causative agent of tuberculosis (TB) and is a major cause of human morbidity and mortality. Crucially, *Mtb* can persist and replicate within host macrophages (MФ) and subvert multiple antimicrobial defense mechanisms. How this is achieved is incompletely understood. Previously, we reported that lipids shed by persistent mycobacteria inhibit NPC1, the lysosomal protein deficient in most cases of the rare, inherited lysosomal storage disorder Niemann-Pick disease type C (NPC). Inhibition of NPC1 leads to a drop in lysosomal calcium levels blocking phagosome-lysosome fusion and thereby leads to mycobacterial persistence.

Studies of mycobacterial lineages have identified events during mycobacterial evolution that result in the acquisition of persistence. We speculated that the production of specific cell wall lipid(s) capable of inhibiting NPC1 activities could have been a critical step in the evolution of pathogenicity. In this study, we have therefore investigated whether lipid extracts from clinical *Mtb* strains representative of multiple *Mtb* lineages, members of the *Mtb* complex (MTBC) and selected non-tubercular mycobacteria (NTM) inhibit the NPC pathway. We have found that the ability to inhibit the NPC pathway was present in all clinical isolates studied from *Mtb* lineages 1, 2 and 4. We also found that lipids from MTBC member, *Mycobacterium bovis* and the NTM, *Mycobacterium abscessus* and *Mycobacterium avium* also inhibited the NPC pathway. However, when lipids were assayed from *Mycobacterium canettii (M. canettii)*, a smooth tubercle mycobacterium, which is considered to resemble the common ancestor of the MTBC no inhibition of the NPC1 pathway was detected. We therefore conclude that the evolution of mycobacterial cell wall lipids that inhibit the NPC pathway evolved early and post divergence from *M. canettii* related mycobacteria and NPC1 inhibition significantly contributes to the ability of these pathogens to persist and cause disease.

**Authors summary:** *Mycobacterium tuberculosis (Mtb)* actively modifies the hostile intracellular environment of host cells to create a niche in which it can persist and replicate. How this is accomplished is incompletely understood. We previously reported an unexpected phenotypic similarity between cells infected with pathogenic mycobacteria and those of the rare, neurodegenerative lysosomal storage disorder, Niemann-Pick disease type C (NPC). Mechanistically, we showed that pathogenic mycobacteria shed lipids that inhibit NCP1, the lysosomal membrane protein that is dysfunctional in NPC and thereby establish conditions that favour their survival. In this study, we have investigated the phylogenetic distribution of lipids that inhibit NPC1. We have screened lipid extracts from multiple clinical mycobacterial isolates representing different *Mtb* lineages, other members of the *Mtb* complex and non-tuberculous mycobacteria for their ability to induce NPC phenotypes. We detected activity in all of the groups tested, but not from *Mycobacterium canettii*, which is believed to resemble the ancestral species. These data indicate that targeting of NPC1 to achieve intracellular persistence evolved early in the evolution of the *Mtb* complex, but after divergence from the presumed ancestral species.

## Introduction

*Mycobacterium tuberculosis* (*Mtb*), the causative agent of tuberculosis (TB) is a major human pathogen [1, 2]. Whilst emergence of multi-drug-resistant strains has added to the challenges of treatment, the ability of *Mtb* to survive and replicate intracellularly in host macrophages (MФ) is key to its success [3, 4]. Typically, bacilli within aerosol droplets are inhaled into the lower pulmonary environment and initially infect resident alveolar MФ, which induces the recruitment of additional myeloid cells leading to the formation of granulomas. Granulomas facilitate infection of other cell types and promote mycobacterial growth and transmission to the next host [3, 5]. This long term intracellular persistence is the basis for latency, which provides a pathogen reservoir that can subsequently become re-activated and promote transmission to a new host [6]. Mycobacteria are phagocytosed following an initial interaction with MФ plasma membrane receptors [7, 8]. However, in contrast to the fate of other ingested particles, which involves phagosome formation and subsequent phagosome-lysosome fusion, *Mtb* and other persistent mycobacteria actively block this process and therefore survive indefinitely within the MФ [9] [5]. Although multiple mechanisms have been proposed to explain the ability of *Mtb* to prevent the fusion of the lysosome with the phagosomes [10],[11] current understanding remains incomplete. The possibility of interfering with the *Mtb*-mediated block in phagosome-lysosome fusion is an attractive target for host-directed *Mtb* therapy.

The mycobacterial cell wall is a complex structure that is composed of multiple carbohydrate and lipid species [12, 13]. Importantly, it represents the interface between the microbe and host cell and therefore plays important roles in determining the outcome of infection. Microbe survival and growth and induction of protective host responses and mycobacterial mechanisms to subvert immunity are dependent upon the architecture and composition of the mycobacterial envelope, [14, 15] [16].

As well as the block in phagosome-lysosome fusion, cells harboring intracellular mycobacteria display several additional characteristics [5] and we have previously reported the unexpected similarity between the cellular phenotypes of *Mtb*-infected MФ and those of the rare, inherited lysosomal storage disease, Niemann-Pick disease type C (NPC) [17]. NPC is caused by mutations in either of two genes, *NPC1* (95% of clinical cases) or *NPC2*. They work cooperatively in a pathway that is involved in lipid trafficking and lysosome: ER contact site formation [18, 19]. In brief, we previously found that all of the cellular phenotypes that define NPC, including a reduction in the calcium content of lysosomes, prevention of phagosome-lysosome fusion, accumulation of specific lipid species (including cholesterol, sphingomyelin and glycosphingolipids (GSLs)) within the endo-lysosomal system were induced in cells infected with pathogenic mycobacteria, but not the environmental non-persistent mycobacterium *Mycobacteria smegmatis*. Very significantly, it was the lipid fraction of the cell wall from persistent mycobacteria that inhibited the NPC pathway. These data are consistent with persistent mycobacteria shedding lipids that inhibit the activity of NPC1, and thereby prevent normal phagosome-lysosome maturation leading to long-term survival [17]. We hypothesized that the evolution of lipids that inhibit NPC1 would corelate with the acquisition of the ability to persist and cause disease. In this study we have therefore assayed the capacity of cell wall lipid extracts to induce NPC cellular phenotypes in RAW 264.7 MФ. We report that extracts from diverse pathogenic mycobacterial species representing the main *Mtb* lineages differentially induce NPC phenotypes with the notable exception of *Mycobacterium canettii (M. canettii)*, which is believed to resemble the putative common ancestor. These findings suggest that the evolution of lipids capable of inhibiting NPC1 evolved early, post-divergence from *M canettii* and correlates with virulence and persistence and therefore represents a new target for therapy.

## Results

### Cell wall lipid fractions from *Mtb* clinical isolates, specific NTM, and *M. bovis* but not *M. canettii*, increase Lystracker staining and granularity of RAW 264.7 MФ

We have reported previously that the volume of the acidic compartment (late endosome/lysosome – LE/Lys) is significantly expanded in genetic NPC1 deficient cells [20] and in RAW 264.7 MФ treated with the cationic amphiphile U18666A, which binds to NPC1 and inhibits its activities [21] or infected with *Bacillus Calmette-Guerin* [17]. Using the fluorescent probe LysoTracker, which selectively accumulates in the acidic compartment, we measured by FACS the relative LE/Lys volume in cells that had been incubated with lipids at a concentration of 100 μg/ml for 48h. We assayed lipids extracts prepared from mycobacterial clinical strains representative of the major *Mtb* lineages (Table 1) and other mycobacterial species (Fig 1). Their activities were compared with vehicle-treated (negative control) and U18666A-incubated (positive control) RAW 264.7 MФ (FACS gating strategy; Supplementary Fig 1). With the exception of strains 232 and 346 from lineage 1 and 212 and 374 from lineage 2 all of the others strains tested induced a statistically significant increase in LE/Lys volume as shown by greater Lysotracker staining (Fig 2A). Strains 119, 173, 293, 367,440, 369 and H37RV increased Lysotracker staining to a level that was not significantly lower than that of cells treated with the NPC1 inhibitor, U18666A. Analysis of the data for each *Mtb* lineage confirmed there was a significant increase in LysoTracker staining of RAW 264.7 MФ incubated with lipid extracts of lineages 2 and 4, but not with lineage 1. This was most likely because two of the four strains from lineage 1 did not induce a significant increase in staining (Fig 2B).

**Table 1.**
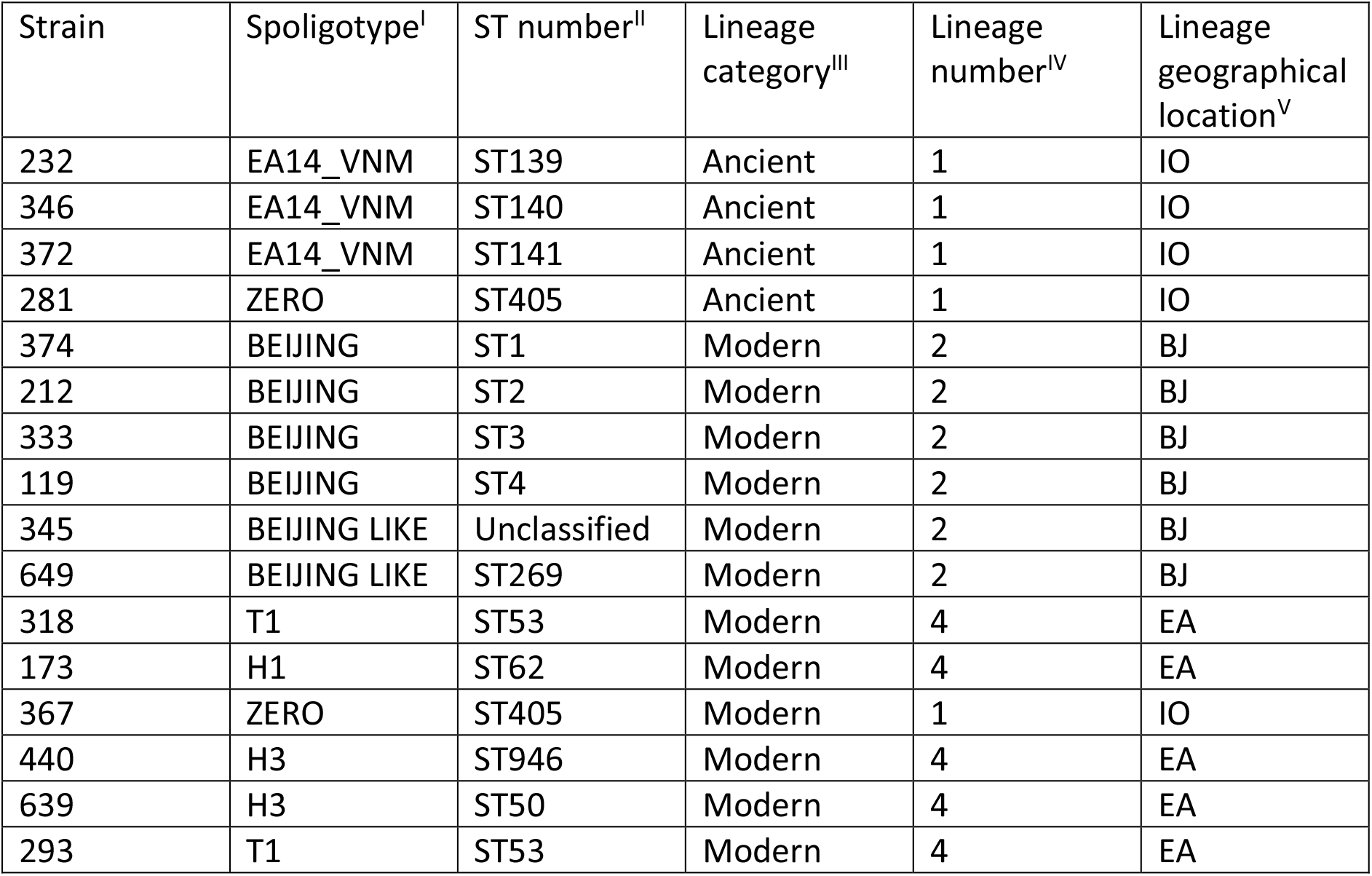
Genotypes of the selected *Mtb* clinical isolates strains used in this study. I and II Classified in accordance with international Spoligotyping database [61]. III Classified based on evolutionary phylogenetic tree [62]. IV Classified in accordance to sequence alignment [34]. V. BJ: East Asian/Beijing; IO: Indo-Oceanic; EA: Euro-American

**Figure 1.**
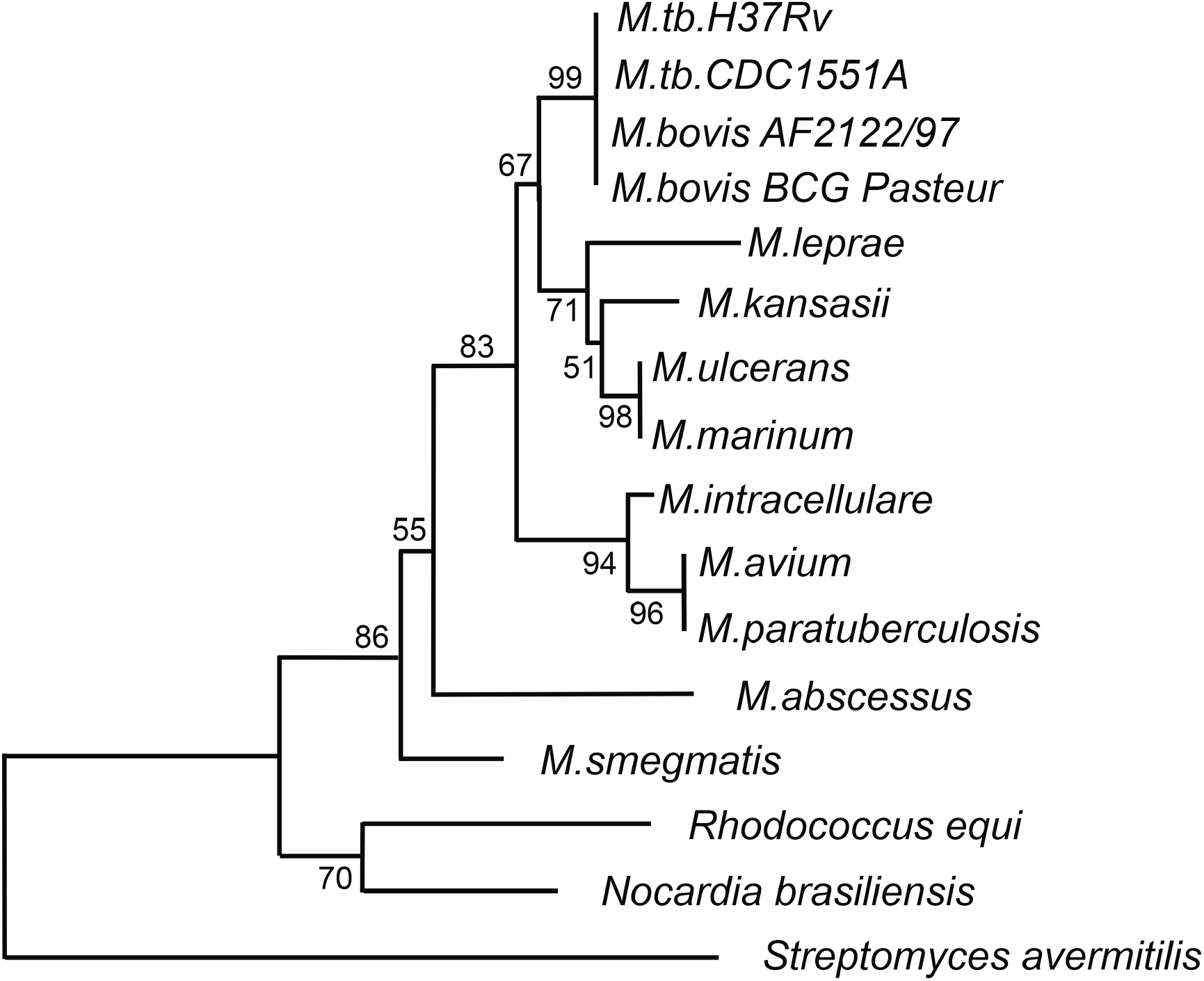
Schematic of phylogenetic relationship of specific mycobacterial species. Phylogenetic tree of mycobacteria based upon the relatedness of the iron-regulated protein HupB [60]

**Figure 2.**
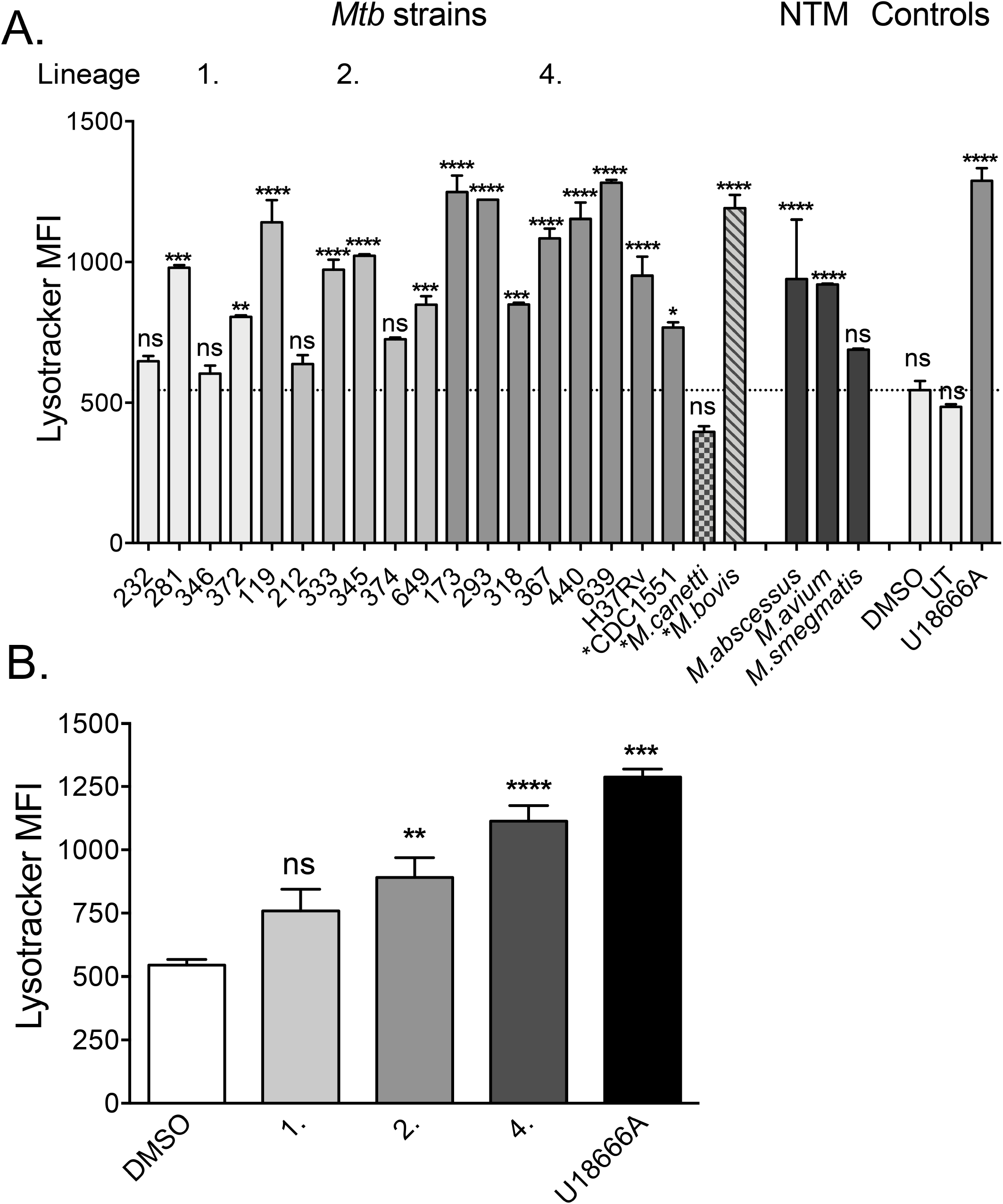
Mycobacterial lipids that can significantly increase Lysotracker staining of RAW 264.7 MФ are widely distributed across *Mtb* clinical strains, members of the *Mtb* complex and non-tuberculous mycobacteria. Panel A. LysoTracker staining of RAW 264.7 MФ treated with 50 μg/ml lipid extract of clinical *Mtb* strains representative of *Mtb* lineages, 1, 2 and 4, *Mtb* complex species and non-tuberculous mycobacteria for 48h. Data is mean ± SEM, N= minimum of 4 replicates per sample. Statistical analysis, 2-way ANOVA, **** *p*<0.0001, *** *p*< 0.001, ** *p*< 0.01. Data is representative of three independent experiments. Panel B. Mean Lysotracker staining of RAW 264.7 MФ treated with lipid from *Mtb* lineages, 1, 2 and 4. Data is mean ± SEM, N= minimum of 16 replicates per lineage. Statistical analysis, 2-way ANOVA, **** *p*<0.0001, *** *p*< 0.001, ** *p*< 0.01. Data is representative of three independent experiments.

Analysis of lipids from other mycobacterial species revealed that the MTBC member *M. bovis* and NTB species *M. abscessus* and *M. avium* also significantly increased LysoTracker staining, whilst extracts from *M. smegmatis* did not achieve significance and *M. canettii* diminished reduced LysoTracker staining relative to control cells (Fig 2A).

We assayed mycobacterial lipid extracts that were isolated at two separate laboratories; one from Imperial College, London and the second from BEI Resources, Virginia. In order to exclude the possibility of a significant variation in activity due to methodological differences in their isolation, we compared lipid extracts of H37Rv from each source within the same experimental assay. The two extracts significantly increased LysoTracker staining to the same extent, which was equal to or great than that of U18666A-treated MФ(Supplementary Fig 2A)

### Expression of LAMP-1 is not increased by exposure to *Mtb* lipids

We imaged RAW 264.7 MФ that had been treated with U18666A or lipid from specific *Mtb* strains and stained for the lysosomal protein LAMP-1. Confocal microscopy suggested that LAMP-1 immunoreactivity was higher in lipid-treated cells (Fig 3A) as compared to controls; however, Q-PCR analysis (Fig 3B) and western blotting (Fig 3C) did not confirm significant increases in either transcription or protein abundance, implying that the differences in confocal images likely reflect a change to protein distribution affecting immunoreactivity.

**Figure 3.**
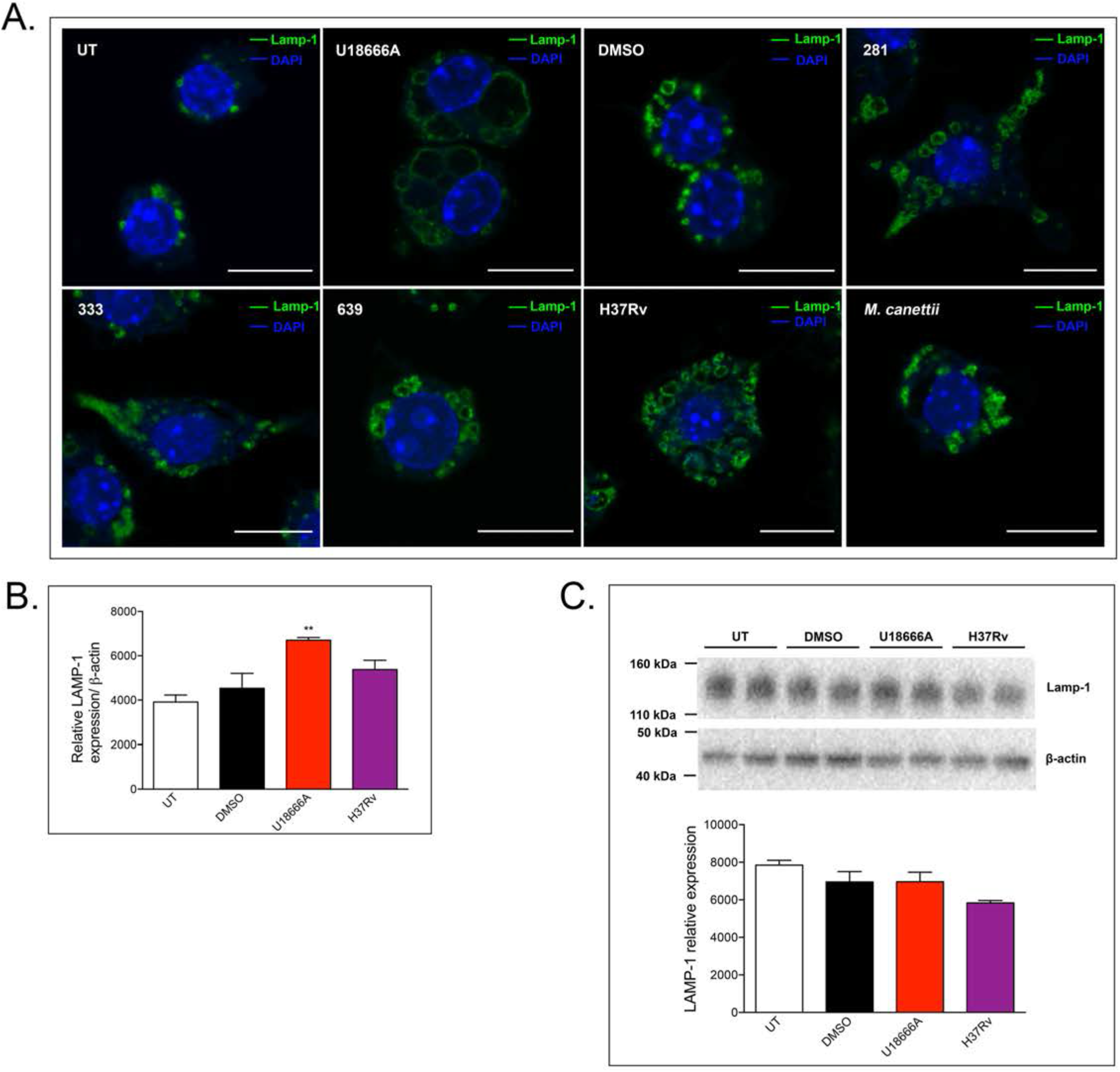
Expansion of the lysosomal compartment in RAW 264.7 MФ in response to treatment with mycobacterial lipid (50 μg/ml for 48h) visualized with anti-LAMP-1 staining. Panel A. Confocal microscopy images of RAW 264.7 MФ stained with anti-LAMP-1 antibody (green). Nucleus is stained with DAPI (blue). Scale bar represents 10 μm. Panel B. Q-PCR of relative LAMP-1 expression in RAW 264.7 MФ, normalized to β-actin. Data is mean ± SEM, N= minimum of 5 replicates per sample. 2-way ANOVA ** *p*< 0.01.Panel C. Western blot of LAMP-1 protein expression in treated RAW 264.7 MФ. Data is mean ± SEM, N= 3 replicates per sample. 2-way ANOVA. No significant differences.

### Cell wall lipid extracts from multiple *Mtb* strains and other mycobacterial species trigger cholesterol accumulation in the endo-lysosomal system of RAW 264.7 M Φ

Lysosomal storage of cholesterol is a hallmark of NPC and was historically used as a method of diagnosis [19]. TB is a pathogen that manipulates host cell lipid metabolism and transforms infected cells into lipid-laden, foamy MΦ that are enriched in cholesterol and indeed cholesterol is important for the formation of granulomas [5, 16, 22]. Cholesterol has also been shown to be a critical energy source for *Mtb* and is essential for persistence [23]. We therefore investigated the capacity of mycobacterial lipid extracts to cause cholesterol accumulation. We selected strains that had significantly enhanced LysoTracker staining (Fig 1A) and were representative of each lineage (lineage 1, strain 281; lineage 2, strain 333; lineage 4, strains 639 and H37Rv) for analysis. In addition, we tested lipid extract of *M. canettii* which did not induce an increase in LysoTracker staining. Evidence of cholesterol accumulation was investigated by filipin staining of fixed cells and biochemical measurement of cell extracts using an Amplex Red assay. As expected, imaging of cells that had been treated with the NPC1 inhibitor U18666A confirmed significant lysosomal accumulation of filipin-positive puncta (Fig 4Ai, which were seen in almost all MΦ (Fig 4Aii). Comparable analysis of cells that had been treated with lipid extracts of the clinical strains 281, 333, 639 and H37Rv also revealed lysosomal puncta (Fig 4Bi), the frequency of which were statistically greater than control (Fig 4Bii). In contrast, relatively few puncta were detected in RAW 264.7 cells treated with an extract of *M. canettii* (Fig 4Bi) and were not significantly greater in number than control (Fig 4Bii). We measured the total cholesterol content of treated cells. Cells treated with U18666A had a statistically greater cholesterol content than control (Fig 4Ci) but although there was a trend of increased cholesterol levels with Mtb lipid treatment, it did not reach statistical significance for any of the strains (Fig 4Cii).

**Figure 4.**
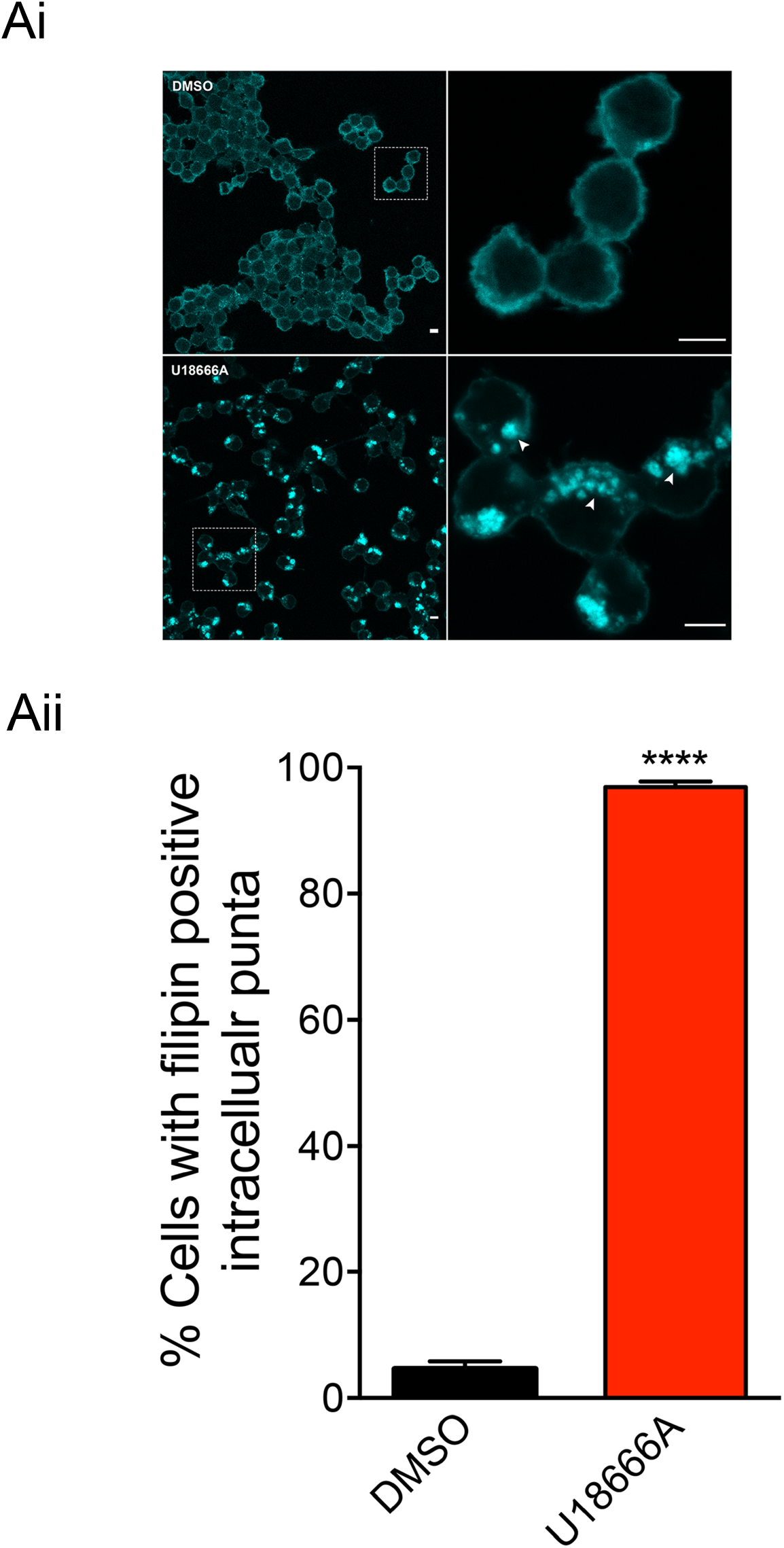
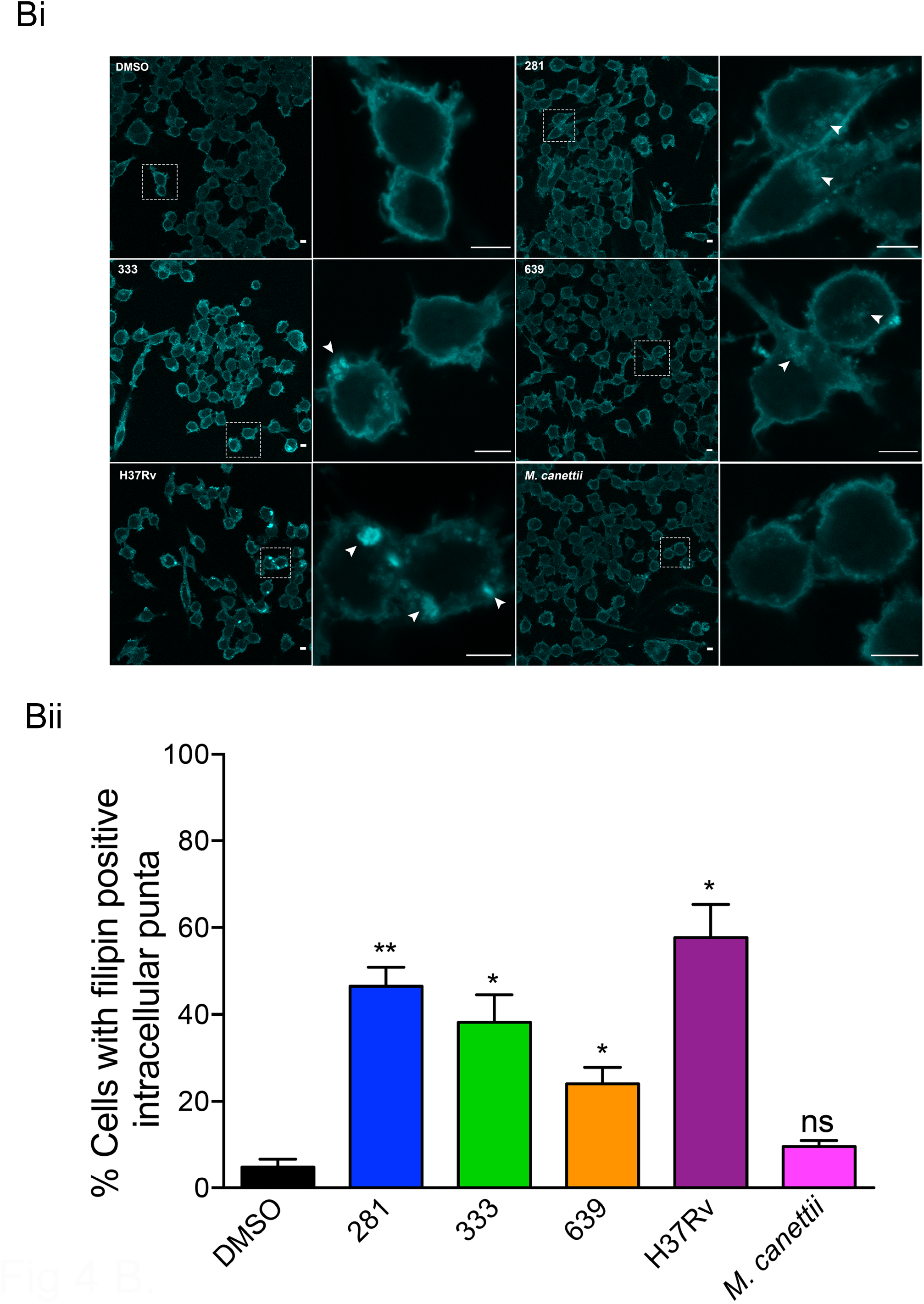
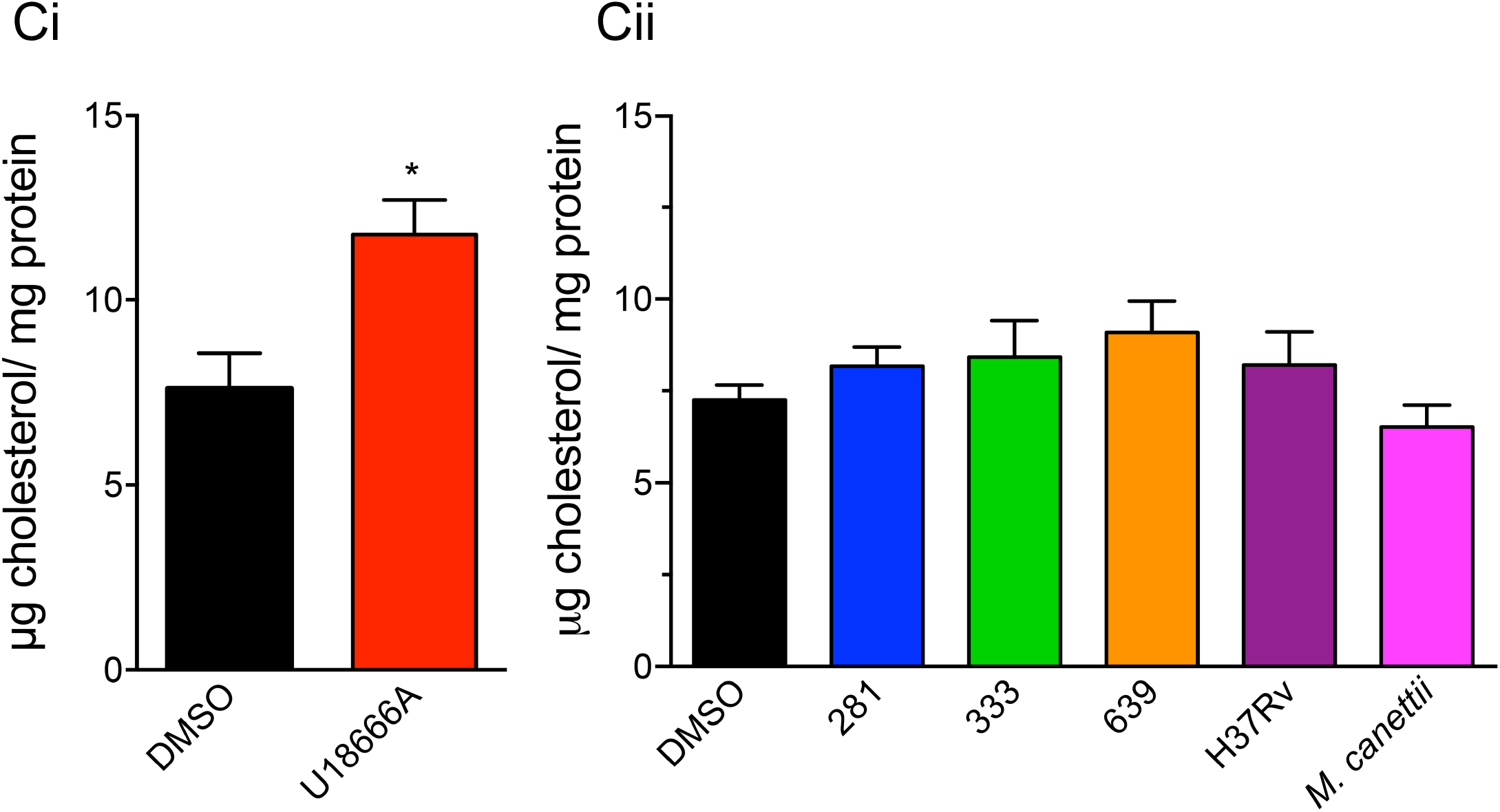
*Mtb* lipids induce a partial redistribution of cholesterol to intracellular puncta in RAW 264.7 MФ. Panel Ai. Confocal microscopy images of vehicle (DMSO) or U18666A treated RAW 264.7 Ф stained with filipin. White arrow heads indicate puncta. Scale bar represents 5 μm. Panel Aii. Quantitation of the frequency of cells with filipin positive intracellular puncta. Data is mean ± SEM, N= minimum of 200 cells per sample. Student t-test **** *p*<0.0001. Panel Bi Confocal microscopy images of RAW 264.7 MФ treated with vehicle (DMSO), *Mtb* strains or *M. canettii* lipids stained with filipin. White arrow heads indicate puncta. Scale bar represents 5 μm. Panel Bii. Quantitation of the frequency of cells with filipin positive intracellular puncta. Data is mean ± SEM, N= minimum of 200 cells per sample. Student t-test. ** *p*< 0.01, * *p*<0.05. ns= not significant. Panel Ci. Amplex red quantitation of cholesterol content of RAW 264.7 MФ treated with vehicle (DMSO) or U18666A. Data is mean ± SEM, N= minimum of 5 replicates per sample. Student t-test. * *p*<0.05. Panel Cii. Amplex red quantitation of cholesterol content of RAW 264.7 MФ treated with vehicle (DMSO) or mycobacterial lipids.

### Exposure to mycobacterial lipids perturb GM1 ganglioside intracellular trafficking

The recycling of glycosphingolipids (GSLs) is perturbed in lysosomal storage diseases, including NPC, resulting in the miss-trafficking of GSLs away from recycling through the Golgi apparatus and instead targets them to the lysosome. This change in lipid trafficking can be visualized by pulse-chase labeling with fluorescently-tagged cholera toxin subunit B that binds to GM1 ganglioside at the cell surface [24]. We used this methodology to assess the extent of altered GM1 trafficking in RAW 264.7 MΦ treated with the different mycobacterial lipids. Confocal microscopy of cells labelled with fluorescently tagged cholera toxin subunit B to monitor trafficking of GM1 ganglioside showed that exposure to U18666A resulted in significant accumulation in the endo-lysosomal compartment (Fig 5A and 5Bi). Similarly, cells treated with lipid from each of the four *Mtb* strains also had significant redistribution of the ganglioside to the lysosome (Fig 5A and 5ii), whereas the lysosomal content of MФ that been exposed to lipids from *M. canettii* was not significantly different from control cells (Fig 5Aand 5Bii).

**Figure 5.**
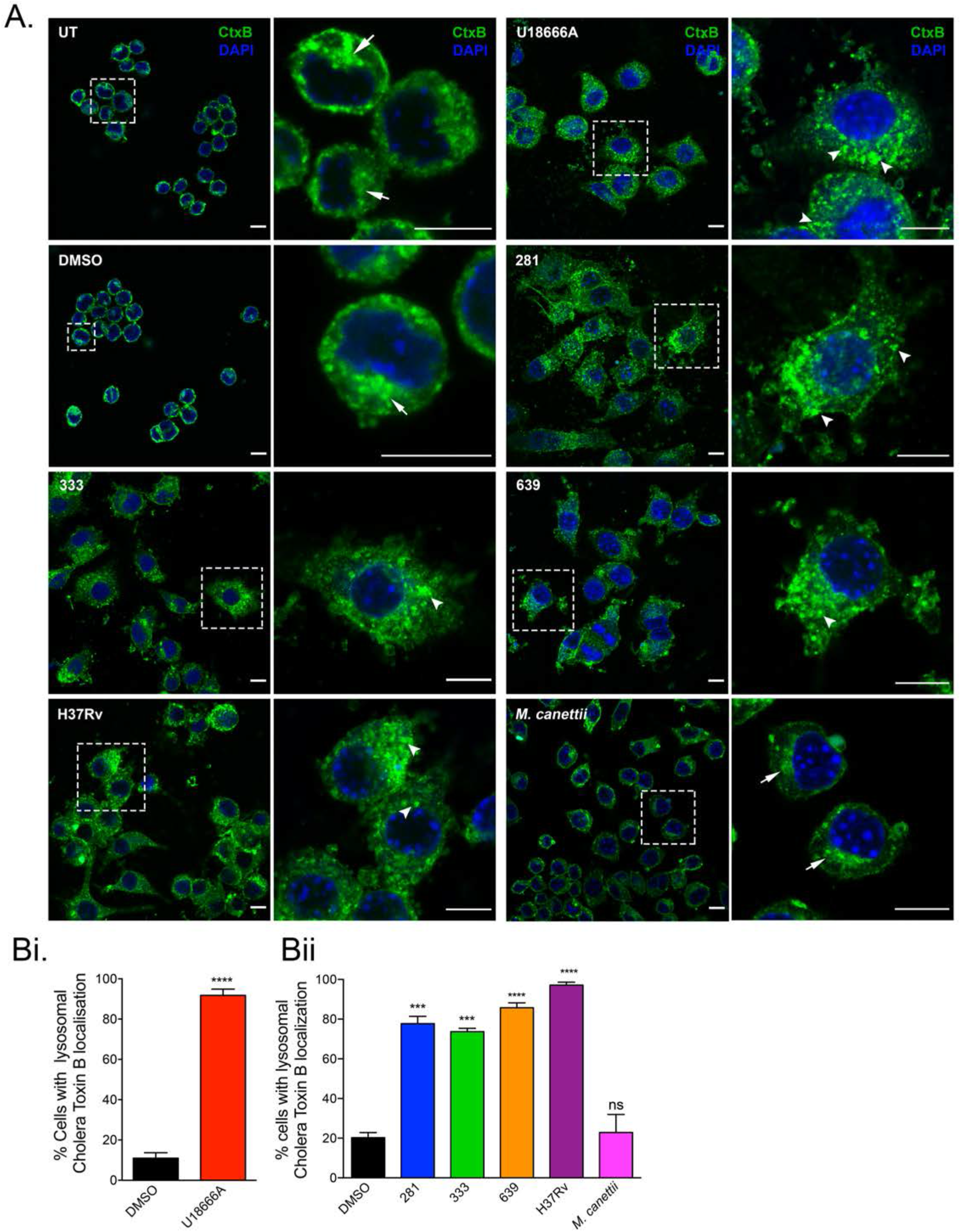
*Mtb* lipids affect GM1 trafficking in RAW 264.7 MФ. Panel A. Confocal microscopy images of RAW 264.7 MФ treated with vehicle (DMSO), U18666A or mycobacterial lipids and pulse-chased with cholera toxin B subunit (green). Nucleus is stained with DAPI (blue). Untreated, vehicle and *M. canettii* lipid treated cells show a predominantly Golgi pattern of staining, indicated by the white arrows. U18666A and lipids from *Mtb* strains induce a punctate pattern, consistent with lysosomal distribution (white arrow heads). Scale bar represents 10 μm. Panel Bi. Percentage of vehicle and U18666A treated cells with lysosomal cholera toxin B localization. Data is mean ± SEM, N= minimum of 200 cells per sample. Student t-test. **** *p*< 0.0001. Panel Bii. Percentage of vehicle and mycobacterial lipid treated cells with lysosomal cholera toxin B localization. Data is mean ± SEM, N= minimum of 200 cells per sample. Student t-test. **** *p*< 0.0001, *** *p*< 0.001. ns= not significant.

### GSL species accumulate in RAW 264.7 MФ incubated with mycobacterial lipid extracts

Impaired lipid transport and catabolism causes the accumulation of multiple different GSLs in NPC cells [25, 26]. We extracted GSLs from RAW 264.7 MФ that had been incubated with cell wall lipid extracts from different mycobacterial species and determined their abundance by normal phase HPLC [27]. HPLC profiles representative of U18666A-treated and DMSO control (Fig 6A) and lipid-incubated MФ extracts and DMSO control are shown in Fig 6B. As expected, U18666A significantly increased total GSL content (Fig 6C), and each of the individual lipid species that we could resolve. Lipid extracts from all four *Mtb* strains were able to significantly increase total GSL levels in treated MФ, but in contrast, total GSLs were significantly lower in cells incubated with *M. canettii* lipids (Fig 6D). Strain 281 from *Mtb* lineage 1 significantly increased eight of the lipid species, lineage 2 strain 33 increased five lipid species, and lineage 4 strains 639 and H37Rv increased four and seven distinct lipid species respectively (Fig 6C and Table 2). *M. canettii* cell wall lipid extract only significantly enhanced the abundance of two specific lipids, albeit to a relatively small extent, and significantly reduced the level of four lipids (Table 2).

**Table 2.** Accumulation of total and specific GSL species in RAW 264.7 MФ treated with, U18666A, lipid extracts from different *Mtb* clinical strains and *M. canettii*. Summary of relative abundance of total GSL content and individual lipid species (indicated by green filled boxes) of RAW 264.7 MФ treated with lipid extract from different *Mtb* strains or *M. canettii* (indicated by blue filled boxes). Data represent mean fold change in lipid abundance relative to DMSO treated control cells. 2-way ANOVA, N= minimum of 5 samples per treatment. **** *p*<0.0001 *** *p*< 0.001 (), ** *p*<0.01, * *p*<0.05, ns = not significant.

| Lipid Extract | U18666A | 281 | 333 | 639 | H37Rv | *M. canettii* |
| --- | --- | --- | --- | --- | --- | --- |
| Lac | ****<br>5.06 | *<br>1.64 | ns<br>1.05 | ns<br>1.17 | *<br>1.90 | **<br>0.60 |
| GA2 | ****<br>6.53 | **<br>1.62 | ns<br>0.93 | ns<br>1.17 | ns<br>1.23 | ns<br>0.88 |
| Gb3 | ***<br>2.96 | ns<br>1.62 | ns<br>1.42 | ns<br>1.51 | *<br>2.40 | ns<br>1.01 |
| GM3 | ****<br>24.68 | ****<br>2.82 | *<br>2.02 | ns<br>2.08 | **<br>5.21 | ***<br>0.62 |
| GM2 | ****<br>36.07 | ****<br>2.82 | ***<br>2.02 | ***<br>2.08 | ****<br>5.21 | **<br>1.20 |
| GA1 | ****<br>4.60 | ****<br>2.24 | ****<br>1.88 | ****<br>1.82 | ****<br>2.23 | *<br>1.23 |
| GM1a | ***<br>1.91 | ****<br>2.02 | ***<br>1.81 | ****<br>1.75 | ****<br>1.56 | ns<br>1.02 |
| GM1b | **<br>1.61 | ***<br>1.55 | **<br>1.41 | ***<br>1.30 | ***<br>1.31 | **<br>0.79 |
| GD1a | ns<br>1.12 | *<br>1.18 | ns<br>1.17 | ns<br>1.06 | ns<br>0.95 | **<br>0.62 |
| Total GSL | ***<br>2.14 | ***<br>1.56 | **<br>1.44 | ***<br>1.34 | **<br>1.28 | **<br>0.79 |

**Figure 6.**
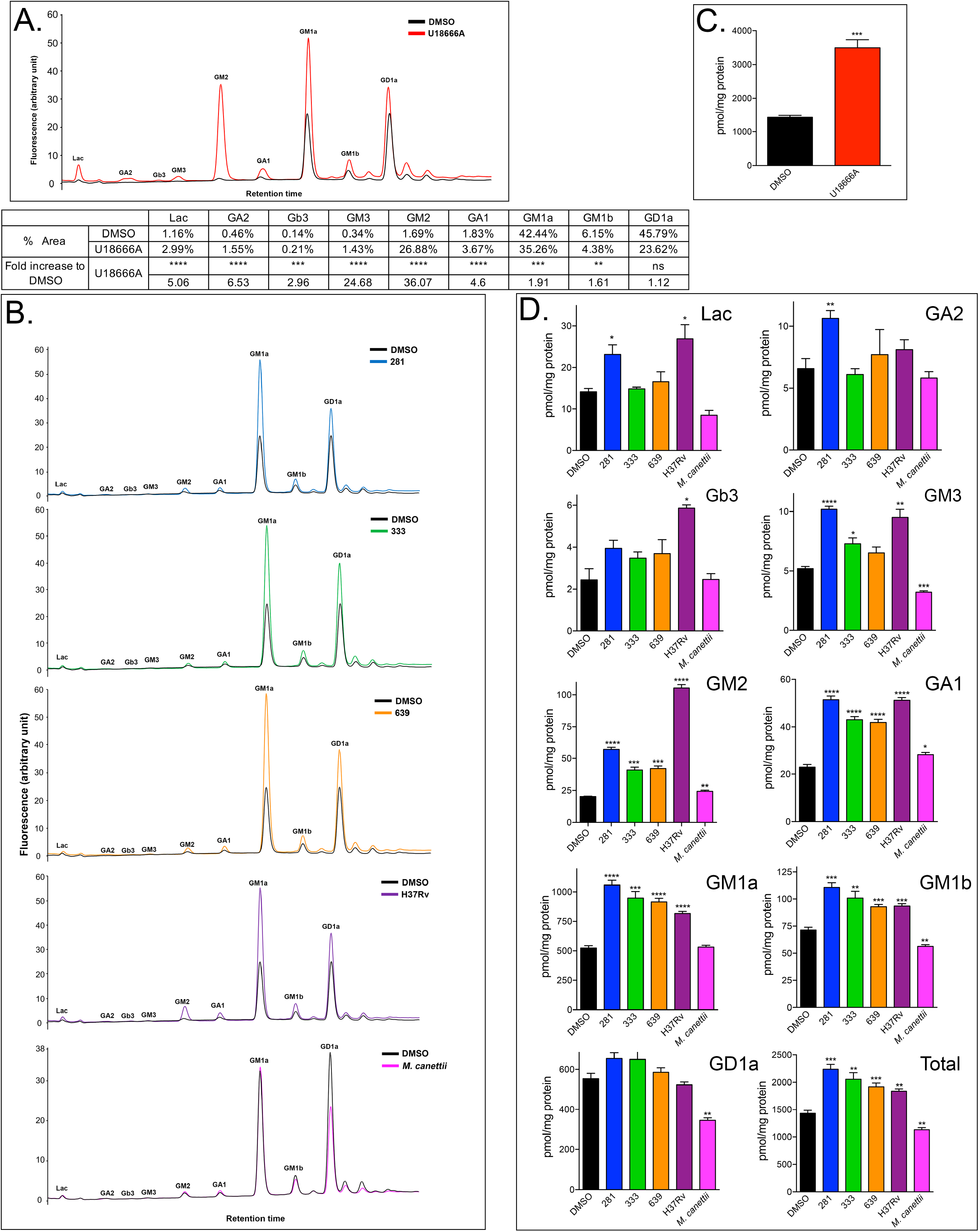
*Mtb* lipids, but not *M. canettii* lipids cause GSL accumulation in RAW 264.7 MФ. Panel A Representative HPLC trace of GSLs from vehicle (DMSO) and U18666A treated cells are overlaid with individual GSL peaks annotated. The table summarizes the percent area of each peak and the fold increase induced by U18666A. Panel B. Overlays of representative HPLC traces of vehicle (DMSO) and individual *Mtb* strains of *M. canettii*. Panel C. Quantitation of total GSLs in DMSO or U18666A treated cells. Data is mean + SEM, N= minimum of 5 replicates per sample. Student t-test. *** *p*< 0.001. Panel D. Quantitation of individual GSL species and total GSLs in DMSO or mycobacterial lipid treated cells. Data are mean ± SEM, N= minimum of 5 replicates per sample. 2-way ANOVA. **** *p*<0.0001, *** *p*< 0.001, ** *p*< 0.01, * *p*<0.05.

### Lipid composition of clinical strains representative of *Mtb* lineages

Using a combination of different solvent conditions and thin layer chromatography (TLC), together with MALDI-ToF mass spectrometry (MS) in the positive and negative ion mode, lipid preparations from each of the *Mtb* clinical strains were analyzed. As shown in Supplementary Fig.3, TLC of extractible lipids from the 17 strains produced qualitatively comparable profiles, but no clear differences in composition were apparent.

## Discussion

TB is caused by closely related members of the MTBC family, which have evolved to become obligate intracellular pathogens. Although individual MTBC species have distinct host preferences, all have adopted a strategy of intracellular persistence. The mechanism through which pathogenic mycobacteria persist within host cells following their ingestion is incompletely understood but is important, as it is a major factor contributing to virulence and is a potential target for host targeted anti-microbial strategies.

In this study, we analyzed the ability of lipid extracts from distinct *Mtb* strains, other MTBC species and NTM to inhibit the NPC pathway in host cells. We have found that this activity is widespread amongst the pathogenic mycobacteria examined but was notably absent from *M. canettii*, a smooth bacillus that belongs to a lineage of progenitor species from which it is believed that the MTBC emerged [28-30]. We hypothesized that the ability of mycobacteria to inhibit NPC1 will correlate with their ability to establish intracellular persistence and be an important event during the evolution of pathogenicity.

We have previous reported evidence revealing the unexpected phenocopying of the complex cellular phenotypes that occur in the lysosomal storage disorder NPC (genetically deficient in NPC1) by MФ infected with pathogenic mycobacteria [17]. Furthermore, infected cell cultures demonstrated that induction of NPC phenotypes occurred not only in cells harboring mycobacteria but also in bystander non-infected MФ, indicating that cell wall-derived lipids released from infected cells and endocytosed by neighboring MФ have identical effects to infection with the live mycobacterium [17].

Whole genome phylogenic studies encompassing the co-evolution of *Mtb* and humans has identified seven prominent lineages of human-adapted mycobacteria that correlate with spread and geographical distribution [31]. We sampled strains representative of three of the major human geographical lineages[32]: lineage 1, East Africa, Philippines and the rim of Indian Ocean; lineage 2, East Asia; lineage 4, modern MTBC of Europe, America and Africa. Lineage 1 is considered representative of ancient human-adapted species, whereas lineages 2 and 4 fall within the classification of modern lineages, based upon the presence or absence of the TbD1 genomic region [33]. Cell wall-derived lipids from twenty strains were chosen to reflect genetic diversity within each lineage [34] and sixteen lipid extracts significantly increased lysosomal volume, consistent with NPC1 inhibition. Of the four strains that were negative, two each were from lineages 1 and 2 and none were from lineage 4. The data revealed differential potencies between the strains that could significantly enhance LysoTracker staining. It was notable that six strains (119, 173, 293, 367, 440 and the laboratory H37Rv) were capable of increasing acidic compartment volume to a level not significantly different from that induced by the pharmacological NPC1 inhibitor U18666A, which is known to bind to and inhibit NPC1 [21] and of these strains, five were from lineage 4, one from lineage 2 and none from lineage 1. However, we cannot at this point absolutely exclude the possibility that the relative effectiveness between cell wall extracts is a result of differential recovery of the active lipid species during the extraction process, rather than the inherent bioactive lipid content of each strain. This issue is likely to be resolved when the identity of the active lipid(s) has been determined and fractionation studies are in progress.

One aim of this study was to determine whether the lipid activity in tuberculous mycobacteria that can induce NPC cellular phenotypes is also present in NTM species and may also be involved in promoting their intracellular survival. *M. abscessus* and *M. avium* are opportunistic pathogens which do not cause TB but can be responsible for severe infections of the respiratory system, skin and mucosa [35-38]. We therefore analysed lipids prepared from other members of the MTBC (*M. bovis* and *M. canettii*) and the atypical or non-tuberculous mycobacteria (NTM), *M. avium* and *M. abscessus*, as well as the environmental non-pathogenic species, *M. smegmatis* (Fig 1). The lack of activity of the lipid extract of the environmental non-persistent species *M. smegmatis* was not unexpected as we have previously shown that infection with live *M. smegmatis* did not cause cholesterol accumulation, GSL miss-trafficking or expansion of lysosomal volume in RAW 264.7 MФ [17].

Significant expansion of LE/Lys was observed in RAW 264.7 MФ exposed to lipid extracts of *M. abscessus* and *M. avium*, indicating that the capacity to induce NPC phenotypes is present within the NTB group of mycobacteria.

The origin and evolutionary history of *Mtb* and the seven other organisms that comprise the MTBC is of particular interest because of the differences in host specificities and pathogenicity, despite their remarkable level of sequence similarity at the nucleotide level [28, 39]. The seven human-adapted lineages are not considered monophyletic. The consensus from multiple genetic studies is that members of the MTBC emerged from progenitor species that likely resembled the smooth tubercule *M. canettii*, which itself diverged prior to the MTBC [40]. It was therefore of significant interest that incubation of RAW 264.7 MФ with the lipid extract from *M. canettii* did not induce any NPC cellular phenotypes, suggesting that the bioactive lipid(s) is either lacking or is of very low abundance in this mycobacterium. Biochemical analysis of mycobacterial cell wall lipids has revealed a discontinuous distribution, consistent with the evolutionary emergence of *Mtb* from an environmental species resembling *Mycobacterium kansasii* via intermediate smooth tubercle bacilli [41]. The diversity of *M. canettii* lipids includes ones that are unique, such as the phenolic glycolipids phenolphthiocerol dimycocerosates that have been lost during the subsequent emergence of MTBC [42, 43]. Molecular and phenotypic studies of the smooth tubercle *M. canettii* have provided evidence for delineating the evolution of *Mtb*, elucidating the underlying mechanisms that were responsible and the development of pathogenicity/persistence and virulence [44, 45]. *M. canettii* strains display significant genome variation [46, 47], which sets them apart from other MTBC. Clinical infection with *M. canettii* is rare and geographically restricted, whilst variants show differential virulence, all of which indicate that it is a far less successful at causing human disease in comparison to *Mtb* [44, 48]. It is therefore possible that the failure of *M. canettii* lipids to inhibit NPC1 indicates use of an alternative mechanism to facilitate infection and intracellular survival that is overtly less effective than the NPC1 inhibition strategy exploited by *Mtb* and other mycobacterial species such as *M. bovis* and *M. avium*.

We also investigated the capacity of mycobacterial lipid extracts to cause lysosomal accumulation of cholesterol in RAW 264.7 MФ, which is a prominent cellular NPC phenotype [19]. The association between cholesterol and *Mtb* infection has been recognized in multiple studies. It is essential for entry of mycobacteria into MФ [49], its utilization is required for persistence [23] and it is important for maintaining dormancy and enabling reactivation [50]. Cholesterol is also used as a critical energy source illustrated by the presence within the mycobacterial genome of genes encoding transport proteins that mediate its import and enzymes necessary for its metabolism [51, 52]. However, to our knowledge the comparative ability of different *Mtb* strains and members of MTBC to manipulate host cell cholesterol has not been explored previously.

Consistent with inhibition of NPC1, lipid extracts from the mycobacteria that affected cholesterol accumulation also caused miss-localization of GM1 ganglioside, which did not occur when RAW 264.7 cells were treated with lipids from *M. canettii*. At this point it is not obvious if pathogenic mycobacteria directly benefit from biochemical changes in the intracellular distribution of host GSLs that result from blocking NPC1-dependent activities and this merits further investigation.

The mycobacterial envelope, which is architecturally and biochemically complex functions as a permeability barrier [13, 53]. Intercalated within the lipid environment of the cell wall are extractable free lipid species that can influence the balance between pathogen and host by affecting pathogenesis and shaping immune responses [12, 54, 55]. In light of the findings that lipid extracts from strains representative of the three *Mtb* lineages increased the relative LE/lys volume to differing extents, we performed lipidomic analyses of sixteen clinical isolates, plus the laboratory strain, H37Rv, to explore lipid diversity and the possibility of correlation between bioactivity and lipid composition. We focused on extractable lipids. As seen on the TLCs, even if qualitatively, no differences were observed, quantitatively, there were some differences in the levels of the respective lipids within and across lineages. These differences could be a factor influencing the phenotypes observed. In addition, it is known that one of the major surface exposed lipids, lipooligosaccharides (LOS), are absent in *Mtb* as opposed to *M. canettii* due to the mutation within the *pks5* locus abrogating synthesis [56]. LOS could therefore be an ideal candidate to explain the suppression of the phenotypes in *M. canettii* compared to *Mtb*, masking more potent surface exposed lipids in the strains used in this study. A similar concept could be applied to *M. smegmatis* which is known to possess a significant amount of surface exposed glycopeptidolipids [57, 58]. It would be of interest in future studies to analyze the lipid extract of *M. canettii* rough morphotype variants I and K, which are deficient for LOS, and test for its capacity to induce NPC1 inhibition. In summary, the findings presented here indicate that the lipid-dependent capacity to trigger specific host cell phenotypes indicative of NPC1 inhibition is broadly distributed amongst pathogenic mycobacteria species and *Mtb* clinical strains. The lack of activity in extracts of *M. canettii* smooth morphotype is consistent with the proposal that the evolution of the lipid(s) that is responsible for NPC1 inhibition occurred after the divergence of MTBC from the ancestral *M. canettii*-like species. Future identification and characterization of the lipid will facilitate a greater mechanistic understanding of how persistent strains establish infection and maintain intracellular persistence.

## Methods

### Mammalian Cell culture

The murine macrophage (MФ) cell line RAW 264.7 was obtained from the ATCC and maintained in RPMI 1640 containing 10% (v/v) fetal bovine serum, 1% L-glutamine and 1% penicillin-streptomycin (all Sigma-Aldrich) at 37°C in 5% CO_2_. Cells were passaged routinely to maintain viability (> 90%).

### Bacterial strains and culture conditions

*M. tuberculosis* strains were cultured in Sauton’s medium as cell surface pellicle and prepared at Imperial College, University of London: *Mtb* clinical strains consisted of 16 clinical strains of *Mtb* collected in Vietnam [34]: 4 Indo-Oceanic (Lineages 1 or named Group I), 6 Beijing (Lineage 2 named Group II) and 6 Euro-American (Lineage 4 or named Group III) to reflect the genotypic diversity found within each lineage in the study Group I (232, 281, 346, 372); Group II (119, 212, 333, 345, 374, 639); Group IIII (173, 293, 318, 367, 440, 639) and *Mtb* H37Rv: non-tubercular mycobacteria: *Mycobacterium abscessus* (ATCC19977) and *Mycobacterium avium* (ATCC 25291) and the environmental species *Mycobacterium smegmatis* mc^2^155 (ATCC 700084). The following lipids were obtained from BEI Resources (Manassas, Virginia, USA): *Mtb* CDC1551 (NR-14838), *Mtb* H37Rv (NR-14837), *M. canettii* (NR-40334) and *Mycobacterium bovis (M. bovis)* (NR-44100).

### Lipid solubilization and addition to RAW 264.7 MФ

Dried lipid extracts were re-constituted at a concentration of 20mg/ ml in DMSO by sonicating for 2 min at 60°C in an ultrasonic water bath. DMSO stocks were diluted to the final concentration in pre-warmed culture medium, sonicated for 2 min at 60°C, allowed to cool and immediately added to cells. RAW 264.7 cells were plated at a density of 5 × 10^4^ cells/ well in 96 well culture plates in complete medium and allowed to adhere overnight. Lipid extracts, 2 μg/ml U18666A (Merck) or vehicle (DMSO only) were added and plates incubated at 37°C, 5% CO_2_ for 48h before analysis.

### Lysotracker Staining and FACS

Lysotracker staining was performed as described previously [20]. In brief, MФ were harvested, washed twice with PBS and stained with 200 nM LysoTracker-green DND-26 (ThermoFisher) for 10 min in the dark. Cells were washed with PBS and re-suspended in buffer containing 5 μg/ml Propidium iodide (Sigma) to allow for exclusion of dead cells and immediately analyzed on a BD FACS-Canto II (Beckton Dickinson). A minimum of 10 000 events were collected for each sample and relative fluorescence values calculated using FlowJo software (Version 10, FlowJo, LLC).

### Filipin staining of cellular cholesterol

Cells were fixed with 4% paraformaldehyde, washed and cellular cholesterol visualized using filipin (Sigma). Cells were incubated with filipin working solution (0.05mg/ml in PBS containing 0.2% Triton X-100) for 1h, washed with PBS and mounted. Imaging was carried out on a Leica-SP8 confocal microscope

### Cholera toxin B subunit staining of GM1 ganglioside trafficking

Cell labelling was performed according to *Chen et al* [24]. RAW 264.7 MФ were washed and then incubated with 0.5 mM Alexa 488 cholera toxin B for 30min, washed, chased in fresh medium for 90min and fixed. Imaging was performed on Leica-SP8 confocal microscope.

### Analysis and quantification of GSLs by Normal Phase High-Performance Liquid Chromatography (NP-HPLC)

GSLs were analyzed essentially as described previously [27]. Aqueous extracts of frozen pellets of control and lipid-treated RAW 264.7 MФ were prepared by three freeze/thaw cycles and extracted with chloroform and methanol overnight at 4°C. GSLs were further purified using solid-phase C18 columns (Telios, Kinesis) and eluted. Fractions were dried down under a stream of nitrogen at 42°C and treated with recombinant ceramide glycanase (rEGCase; GenScript) to release ceramide-linked glucosylceramide derived GSL oligosaccharides. The liberated free glycans were fluorescently labeled at 80°C for 60 min with anthranilic acid (2AA) and sodium cyanoborohydride. Purification of labeled glycans and removal of excess 2AA was achieved by passing reaction mixes over DPA-6S SPE columns (Supelco). 2AA-labeled oligosaccharides were separated and quantified by normal-phase HPLC as described by Neville *et al* [59].The NP-HPLC setup consisted of Waters Alliance 2695 separation module and an in-line Waters 2475 multi l-fluorescence detector set at E x l360nm and Em l425um. The solid phase was a 4.6 × 250mm TSK gel-Amide 80 column (Anachem). Glucose unit values (GUs) were determined using a 2AA-labeled glucose homopolymer ladder (Ludger). Individual GSL species were identified by their GU values and quantified by comparison of integrated peak areas with a known amount of 2AA-labeled BioQuant chitotriose standard (Ludger). Protein content of the cell lysates was determined using BSA assay (Sigma).

### TLC and MALDI lipid fingerprinting

Heat-killed mycobacteria were first washed 3 times with PBS. The pellets were then submitted to CHCl_3_/MeOH 1:2 (v/v) extraction for 12h at room temperature followed by one CHCl_3_/MeOH 1:1 (v/v) extraction and one CHCl_3_/MeOH 2:1 (v/v) extraction for 3h at room temperature. Pools extracts were concentrated and evaporated to dryness. Total lipids extracted were normalized to the dried weight of lipid extract. The dried lipids extracts were resuspended in CHCl_3_ at a final concentration of 20 mg/mL. 5μL, equivalent to 100 μg, were loaded on TLC.TLC were run in four different solvent system Petroleum Ether/Diethyl Ether 95:5 (v/v); CHCl_3_/MeOH 9:1 (v/v); CHCl_3_/MeOH 8:2 (v/v) and CHCl_3_/MeOH/H_2_O 60:25:4 (v/v/v). Then, TLCs were sprayed with a solution of 5% phosphomolybdic acid in 100% ethanol and heated at 200°C for 2 minutes in order to reveal the lipids. All experiments were performed in triplicate for statistical confidence.

For lipid fingerprinting, 5 ml of heat-inactivated mycobacterial culture was re-suspended in 500 μl of distilled water, washed three times with double distilled water and re-suspended in 100 μl of double distilled water. 0.4 μl of the mycobacterial solution was loaded onto the target and immediately overlaid with 0.8 μl of a 2, 5-dihydroxybenzoic acid (DHB) matrix used at a final concentration of 10 mg/ml in chloroform/ methanol (CHCl_3_/MeOH/TFA) 90:10:1 v/v/v. Mycobacterial solution and matrix were mixed directly on the target by pipetting and the mix was dried gently under a stream of air. MALDI-TOF MS analysis was performed on a 4800 Proteomics Analyzer (Applied Biosystems) using the reflectron mode. Samples were analyzed by operating at 20 kV in the positive and negative ion mode using an extraction delay time set at 20 ns.

## Acknowledgements

This study was supported by a Wellcome Trust Investigator in Science award to FP 202834/Z/16/Z. FP is a Wolfson Royal Society Merit Award holder. This study was supported by the MRC Confidence in Concept Fund and the ISSF Wellcome Trust Grant 105603/Z/14/Z (GL-M).

## Figure Legends

**Supplementary Figure 1.**
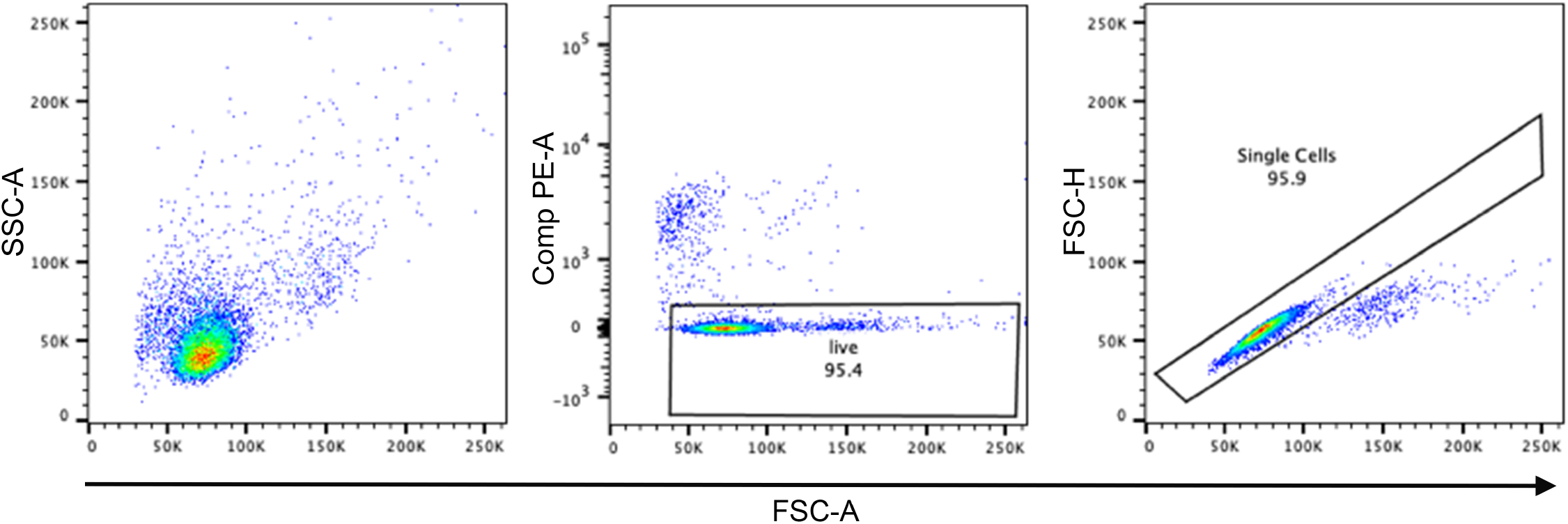
FACS gating strategy for LysoTracker staining of RAW 264.7 MФ.

**Supplementary Figure 2.**
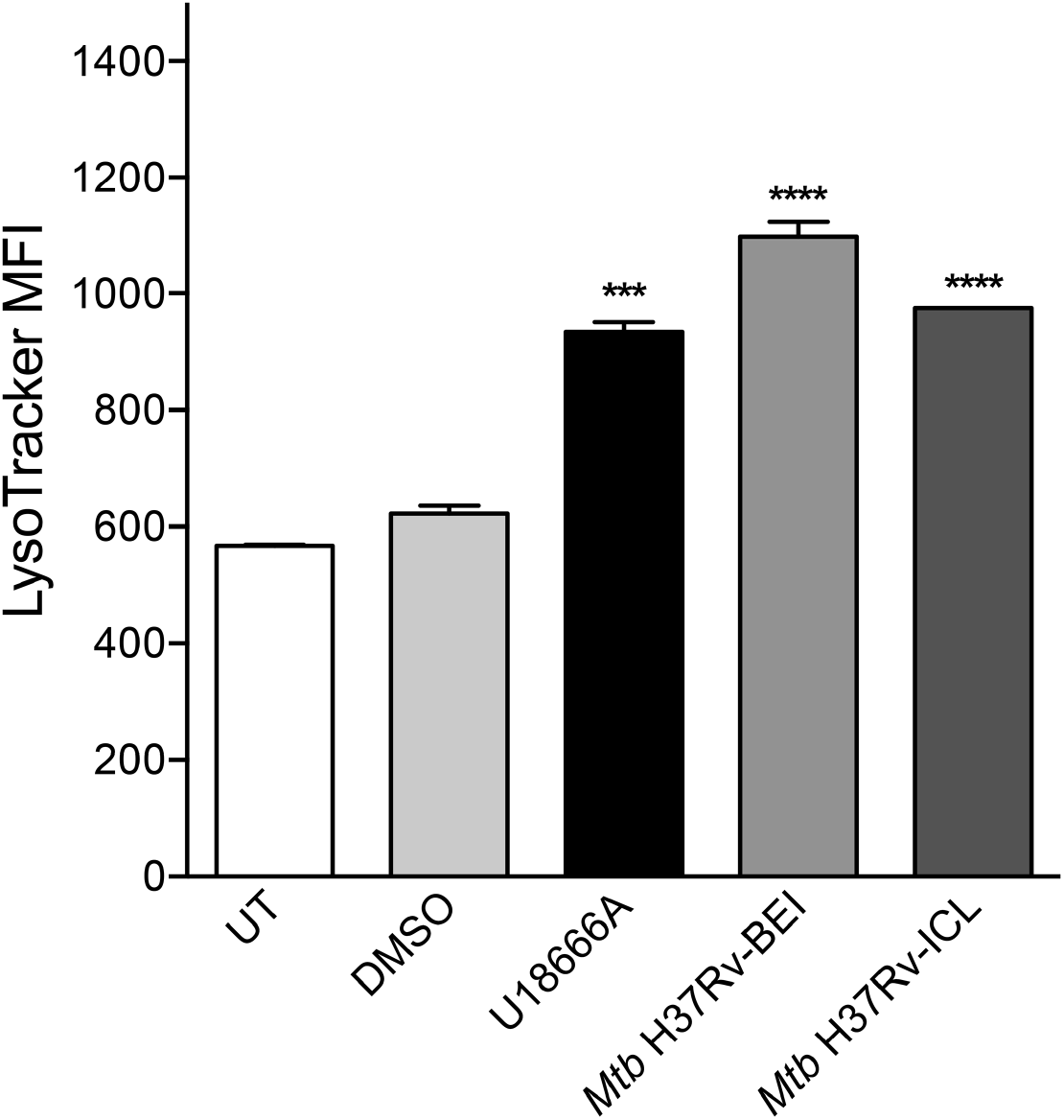
*Mtb* H37Rv lipids from two independent sources increase Lysotracker staining of RAW 264.7 MФ to an equivalent extent and are comparable to U18666A.

**Supplementary Figure 3.**
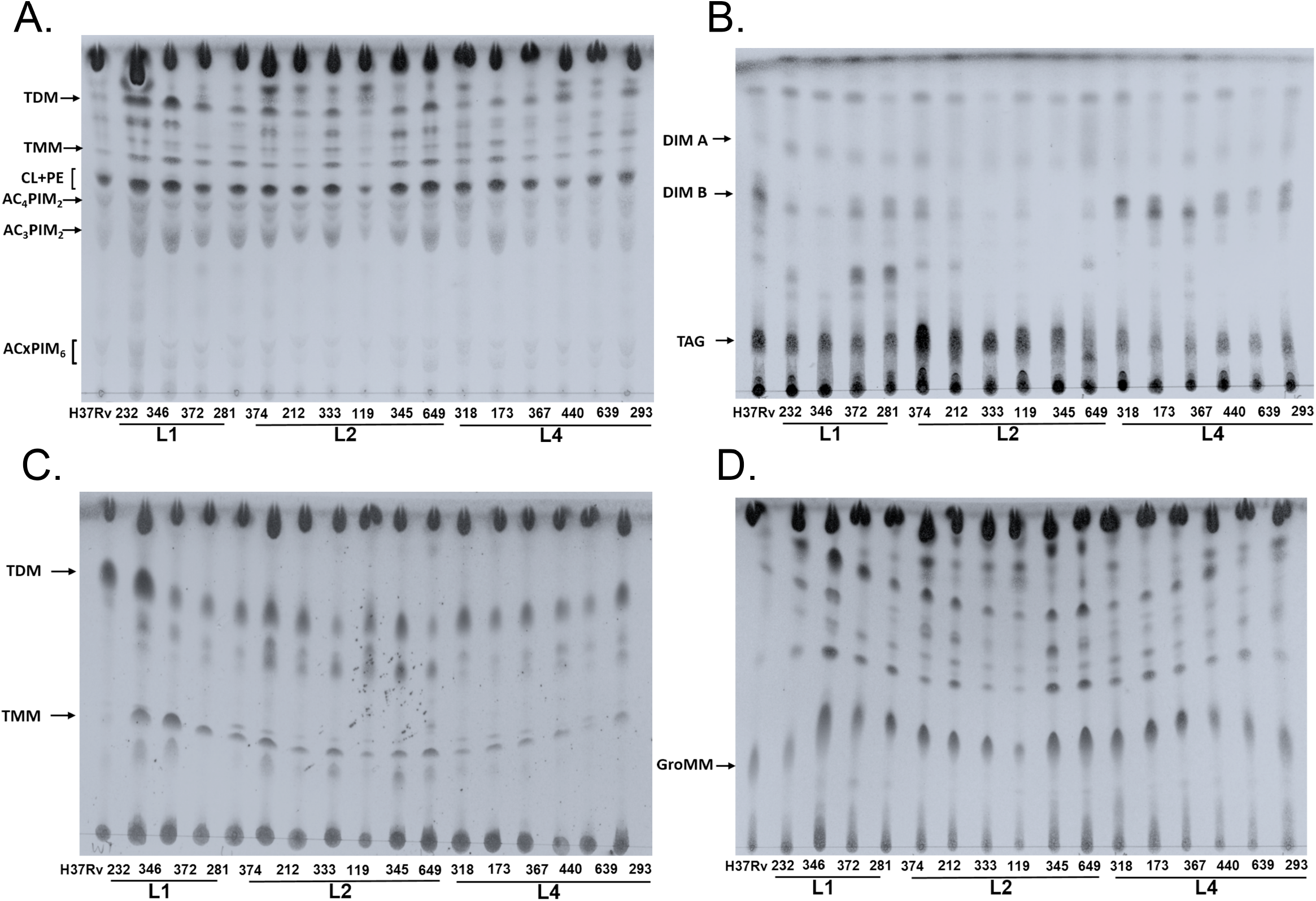
Thin-layer chromatograms of lipids from different *Mtb* strains. TLCs were run in four different solvent systems. Petroleum Ether/Diethyl Ether 95:5 (v/v) (Panel A); CHCl_3_/MeOH 9:1 (v/v) (Panel B); CHCl_3_/MeOH 8:2 (v/v) (Panel C) and CHCl_3_/MeOH/H_2_O 60:25:4 (v/v/v) (Panel D). The migration of specific lipid species is indicated.

